# Screening and identification of two pheromone receptors based on the coadaptation of pheromones and their receptors in rats

**DOI:** 10.1101/2022.12.27.522049

**Authors:** Wei-Chao Wang, Yao-Hua Zhang, Guan-Mei Hou, Yan-Yan Sun, Yi-Jun Wu, Jian-Xu Zhang

## Abstract

The coadaptation or genetic coupling of senders and receivers of sex signals in some animals has been described, but no empirical evidence shows whether sex pheromones and their receptors undergo coadaptation in mammals. In this study of the brown rat (*Rattus norvegicus*), we found markedly higher levels of two predominant male pheromones (2-heptanone and MUP13) in the North China subspecies [*R. n. humiliatus* (RNH)] than in the Northeast China subspecies [*R. n. caraco* (RNC)] by gas or liquid chromatography and mass spectrometry. Coincidently, two vomeronasal receptor genes (*Vom1r68* and *Vom2r53*) were found to be expressed at higher levels in RNH females than in RNC females and were thus considered candidate receptors for 2-heptanone and MUP13, respectively. An immunofluorescence analysis showed that these two VR receptors colocalized with mTmG on the membrane of HEK293-T cells. We also verified the responsiveness of Vom1r68 to 2-heptanone and Vom2r53 to MUP13 in HEK293-T cells by calcium imaging. In conclusion, we screened and identified the receptors of two pheromones based on the coadaptation of pheromones and their receptors, which further verified their coevolution.

## Introduction

Chemical communication plays a key role in almost all behavioural aspects in mammals (1–3). Pheromones are chemical communication signals with certain chemical components that are produced in specialized skin glands, urine, and lacrimal glands, among others, and vomeronasal organs (VNOs) are responsible for the specific detection of pheromones (2–5). The screening and identification of vomeronasal receptors for a particular pheromone from hundreds of vomeronasal receptors requires complex and expensive techniques (6, 7). More than a dozen pheromones have been identified from the large number of chemicals, particularly in mice and rats, but only a few pheromone-matched receptors have been identified from approximately 250 V1Rs (VR family 1) and 100 V2Rs (VR family 2) of VNOs (4–14). Based on coevolution or mate choice hypotheses, such as those related to good genes and Fisherian sexy sons, the coevolution or genetic coupling of signals and their receptors can occur (15–19). This phenomenon occurs in many species, as exemplified by acoustic communication in crickets and grasshoppers, visual communication in medaka and chemical communication in corn borer moths (15, 17). In *Drosophila*, the expression of a desaturase gene, *Desat1*, in neural and nonneural tissues mediates the coevolution of sex pheromone perception and emission, respectively (18). Therefore, in murine rodents, we sought to screen pheromone-specific receptor candidates according to their coadaptation with pheromones (20).

In mammals, urine is the most common source of pheromones, and urine marking is the most important method of pheromone transmission (1, 2, 4). Our laboratory recently identified 2-heptanone, 4-heptanone, 9-hydroxy-2-nonanone, MUP13 and OBP3 as male pheromones of rats (10, 14). In rats, MUP13 is an innate pheromone, and 2-heptanone is a learned pheromone (21). Whether innate or learned, pheromones require corresponding receptors to be accepted (1). Male mouse urine clearly contains a markedly higher abundance of volatile pheromones than male rat urine (4, 10, 12). In rat urine, 2-heptanone is the only predominant volatile pheromone, and the major urinary protein (MUP) composition is simpler than that of mouse urine (13, 22–25). Therefore, the rat is presumably a simpler model for studying mammalian pheromone communication systems than the mouse (10, 12–14).

In wild populations of brown rats (*Rattus norvegicus*) (the conspecific ancestor of laboratory rats), two rat subspecies, the North China subspecies *R. n. humiliatus* (RNH) and the Northeast China subspecies *R. n. caraco* (RNC), are scientifically recognized based on distinct morphological differences, and their core populations are separated by 1,000 km (26, 27). We recently used whole-genome analysis to examine the genetic divergence of brown rat populations, including these two rat subspecies (20, 26, 28, 29). The core populations of RNC and RNH are differentiated both genetically (e.g., in olfactory receptor and pheromone-related genes) and phenotypically (e.g., in coat colour, body type, and urine compounds) (20, 28, 29). Additionally, both male pheromones and female receptors are more highly expressed in RNH than in RNC rats, and we thus questioned whether and how male pheromones coadapt with their receptors in female VNOs (20, 26). Two male-predominant pheromones, 2-heptanone and MUP13, exhibit a stable difference between these two rat subspecies, and RNA-seq revealed a few potential V1R and V2R receptors that covary with them (20). Based on the above mentioned coevolution hypothesis and mate choice hypothesis, we speculated the existence of a specific relationship between these pheromones and pheromone receptors that is worth further verification (16, 18).

Mammals exhibit both a main olfactory system and an accessory olfactory system: the former primarily detects general odours in the environment, and the latter perceives most pheromonal signals (1). The accessory olfactory system consists of a pair of VNOs and central projection areas, including an accessory olfactory bulb connected to the amygdala, hypothalamus and cortex (1, 30). Vomeronasal sensory neurons (VSNs) are molecularly and functionally distinct in rodents (31, 32). The VSNs of the apical layer expressing V1R mainly recognize volatile pheromones and sulfated steroids (6, 31–35), whereas the basal layer expressing members of V2R mainly recognizes nonvolatile pheromones, such as MUPs and peptides (7, 8, 32, 34–36). Pheromone-detecting receptor neurons are highly specific and ultrasensitive (1, 33, 37). Therefore, each of two predominant pheromones, 2-heptanone and MUP13, should have specific receptors in the corresponding V1R and V2R families in rats. The identification of pheromone-specific receptors is of great significance for revealing the molecular basis of pheromone perception in mammals (35).

With the discovery of odorant receptor genes in rats, the study of chemical senses entered the molecular age (38–40). However, only a few pheromone receptors have been identified thus far in mice (6, 7, 41). The deorphanized receptors were mostly tested in vitro via expression in heterologous systems (baculoviruses, insect ovarian cells, yeast, *Xenopus* oocytes, and HeLa and HEK cells) (42). A heterologous system allowed the identification of olfactory receptors and vomeronasal receptors for a set of biologically relevant ligands in insects and mice, which made it possible to assess receptor responsiveness to odorants (35, 43–46). Currently, we are using HEK293-T cells as a heterologous system to express pheromone receptor candidate genes and verify the response to certain pheromones via calcium imaging.

By RNA-seq, we recently identified a few vomeronasal receptor genes among approximately 180 vomeronasal receptor genes that showed differential expression between RNH and RNC rats, and the VSNs were more sensitive to 2-heptanone and MUP13 in RNH compared with RNC rats (20). Additionally, most urine-borne volatile compounds and total MUPs were found at increased levels in RNH rats than in RNC rats (20). In the current study, we critically analysed the pheromones and vomeronasal receptor genes showing differences between the RNH and RNC rat subspecies by gas or liquid chromatography and mass spectrometry, RNA-seq, and qPCR and examined their one-to-one correspondence via the transient expression of membrane proteins in HEK293-T cells and calcium imaging approaches.

## Materials and Methods

### Subjects

The rats used in this study were more than ten generations from ancestors captured from wild populations. For breeding, males of the same subspecies were caged in groups of three or four same-sex siblings of the same subspecies after being weaned at 4 weeks of age and caged in plastic rat cages (37 × 26 × 17 cm) (Suzhou Fengshi Laboratory Animal Equipment Co., Ltd., Suzhou, China) with wood shavings for bedding (Beijing Keao Xieli Feed Co., Ltd., Beijing, China). Standard rat chow and tap water were provided ad libitum. The colonies were maintained under a 14-h light:10-h dark cycle (lights on at 19:00) at 23 ± 2 °C. All male rats were 5 to 12 months of age, sexually naïve, and caged individually for 2 weeks prior to initial urine collection. Six male rats of each subspecies served as urine donors (20). All the females were 3-6 months of age and sexually naïve, had a 4- or 5-day oestrous cycle and were killed at oestrus. The phase of the oestrous cycle was determined by vaginal smear cytology, and only females with regular oestrous cycles were used in the study. The animal handling procedure complied with the institutional guidelines for animal use and care established by the Institute of Zoology, Chinese Academy of Sciences. Ethical approval was obtained from the Institutional Ethics Committee of the Institute of Zoology, Chinese Academy of Sciences (20). Our study was performed in accordance with ARRIVE guidelines.

### Urine collection and tissue samples

For urine collection, rats were individually caged in clean metabolic cages for 8 h daily during the dark phase of the light cycle. Standard rat chow and water were provided ad libitum. The urine from each metabolic cage was collected in a tube immersed in an ice box. The urine samples were stored at −20 °C prior to use. The metabolic cages were washed thoroughly with water and sterilized between collections.

The VNOs used for qPCR were dissected from 4 females of each subspecies. We killed the rats via cervical dislocation, opened the mouth with forceps, and visualized the palate. An incision was made in the upper part of the palate. We removed the palate membrane with microdissecting forceps, cut the upper and lower parts of the nasal septum with iris scissors, and then dissected the VNO. The livers used for absolute qPCR quantification were dissected from 6 male rats of each subspecies after the animals were killed via cervical dislocation. All samples were rapidly sampled and preserved in separate cryopreservation tubes immersed in liquid nitrogen prior to use. All procedures were conducted within 10 min after death, and the samples were rapidly frozen with liquid nitrogen.

### Gas chromatography–mass spectrometry (GC–MS) assay of urinary volatile compounds

For the extraction of compounds from urine samples obtained from 22 RNH males and 21 RNC males, we mixed 150 μl of dichloromethane (purity > 99.5% DIMA Technology, Inc.) with 150 μl of urine, stirred the mixture thoroughly, stored it at 4 °C for 12 h, and then used the bottom phase (the dichloromethane layer) for chemical analysis. Chemical analysis was performed with an Agilent Technologies Network 6890N GC system coupled with a 5973 Mass Selective Detector with the NIST/EPA/NIH Mass Spectral Library (2008 version; Agilent Technologies, Inc., USA). The GC–MS conditions and parameters were the same as those described in the literature (12). We injected 4 μl of urine in the splitless mode. Tentative identification was performed by comparing the mass spectra of the GC peaks with those in the MS library (NIST 2008).

Diagnostic fragments at m/z 58 and m/z 71 indicate the presence of ketones. Eight of the tentatively identified compounds, 4-heptanone, 2-heptanone, dimethyl sulfone, 4-methyl phenol,
4-ethyl phenol, and squalene, were further confirmed by matching the retention times and mass spectra with authentic analogues (all purity >98%; ACROS Organics). For a particular compound, the abundance was quantified based on the GC peak area, and the relative abundance was determined as a percentage of the sum of the peak areas from all targeted GC peaks.

### Quantitative real-time PCR (qPCR) of vomeronasal receptor genes and absolute quantitative PCR of hepatic *Mup* genes

The TRIzol method (Life Technologies, USA) was used for the extraction of total RNA from the VNOs of nine females and the livers of six males of either the RNH or RNC subspecies, and HiFiScript gDNA Removal RT Master Mix (CWBIO, China) was employed for the reverse transcription of cDNA according to the manufacturer’s instructions. qPCR was then conducted using the Magic SYBR Mixture (CWBIO, China) and a PIKOREAL 96 qPCR system (Thermo Scientific, USA). Specific oligonucleotide primers were designed using the NCBI-Primer-BLAST online tool (https://www.ncbi.nlm.nih.gov/tools/primer-blast/index.cgi?LINK_-LOC=Blast_Home). GAPDH was used as the housekeeping gene. The oligonucleotide primer sequences are shown in Supplementary Table S1. The 15-μl reaction included 7.5 μl of 2× Magic SYBR Mixture, 0.3 μl of cDNA template, 0.3 μl of each primer (10 μM) and 6.9 μl of RNase-free water. The amplification reaction was performed under the following conditions: denaturation for 30 s at 95 °C and 40 cycles of 95 °C for 5 s and 60 °C for 30 s. The temperature program for the melting curve analysis was 95 °C for 15 s, 60 °C for 1 min, 95 °C for 15 s and 50 °C for 30 s. The VNO qPCR data were analysed using the 2^ΔΔCT^ method (47).

The absolute quantitative PCR analysis of hepatic *Mup* genes was performed as follows: The rat *Mup13, Obp3, Mup4, Mup5, Pgcl2* and *Pgcl3* cDNAs in male rat livers were amplified by PCR (PrimeSTAR DNA Polymerase Takara, Japan). The oligonucleotide primer sequences employed for this purpose are shown in Supplementary Table S1. The target gene was ligated into the pGM-T vector. The constructed plasmid was transformed into DH5α cells (CB101, Tiangen, China). Positive clones were selected and sequenced using universal primers (T7 promoter and SP6 terminator primer). The strains were expanded and cultured with a TIANprep Mini Plasmid Kit (DP103, Tiangen, China). The known standard curves were used to quantify the initial amount of the target template of the unknown sample; 5-fold dilutions of all the standard products were used as the templates, and qPCR was performed. The standard curve was drawn with the logarithm of the initial copy number of the target template as the abscissa and the detected Ct value as the ordinate. Linear regression equations were obtained: slope = 3.100-3.582, amplification efficiency E = 10^(−1/slope)^ −1, and correlation coefficient R^2^ > 0.99. Copies, unless otherwise indicated, refer to abundance of cDNAs derived from 100 ng of RNA, and absolute values were determined by control PCR assays involving plasmid titrations.

### Sodium dodecyl sulfate–polyacrylamide gel electrophoresis (SDS–PAGE) of MUPs

The relative abundances of MUPs in six males of each subspecies were qualified by SDS–PAGE using a Mini-Protean system (Bio-Rad, USA). Eight microlitres of urine from each male was diluted 2.5-fold by mixing with 4 μl of 5× SDS–PAGE loading buffer (Solarbio, Beijing, China) and 8 μl of sterile water. Mixed protein denaturation was performed at 100 °C for 5 min in a Dry Bath Incubator (MIULAB, Hangzhou, China). Four microlitres of each mixed sample and PageRuler™ Prestained Protein Ladder (Thermo Fisher Scientific, USA) was fractionated on 15% SDS–PAGE gels. Protein gels were run at a constant voltage of 150 V for 1 h and coloured using Coomassie bright blue dye. The target band was photographed with a ChemiDoc MP system (Bio-Rad, USA). The relative abundance of MUPs was analysed with the ImageJ program (National Institutes of Health, USA). One sample was selected as the standard control, fractionated on each gel to correct the deviation between runs, and normalized to the relative abundance in all other samples (20).

### In-gel digestion of MUPs

Each selected band was removed from the SDS–PAGE gel, cut into 3-4-mm^3^ pieces and placed into 1.5-ml microcentrifuge tubes. The gel pieces were destained using 50 μl of 50 mM ammonium bicarbonate (NH_4_HCO_3_)/50% (v/v) ACN for 30 min at 37 °C. The gel pieces were incubated with 10 mM dithiothreitol (DTT) for 60 min at 56 °C. The DTT was then discarded, a 55 mM iodoacetamide (IAM) stock solution was added to each tube, and the resulting mixture was incubated for 60 min in the dark. After the IAM was discarded, the gel pieces were washed twice using 200 μl of ACN. The gel pieces were dried with 200 μl of ACN and digested with 10 ng/μl Lys-C (sequencing grade, Promega, USA) in 50 mM NH_4_HCO_3_ overnight at 37 °C (14).

Supernatants were collected in fresh 1.5-ml tubes on the following day, and the remaining peptides were extracted from the gel pieces in 80 μl of a solution comprising 50% ACN, 5% trifluoroacetic acid (TFA) and 45% deionized water via sonication for 15 min. The tubes were briefly centrifuged for 30 s, the supernatants were collected, and the process was repeated. All supernatants from each band were pooled, dried in a Speed-Vac evaporator (Thermo Scientific, USA), and mixed with 12 μl of 0.1% formic acid. Thereafter, the solution of each gel piece was transferred to an ultrafiltration tube (UFC30HV25, Lithuania) and centrifuged at 12800 rpm for 15 min. The final solution of each band was stored at −20 °C until use. Unless stated otherwise, all procedures were performed at room temperature (14).

### Liquid chromatography-tandem mass spectrometry analysis (nLC–MS/MS) of MUPs

The extracted peptides were analysed using an Easy-nLC 1200 system (Thermo Scientific, USA) coupled to an Orbitrap Exploris 480 mass spectrometer through an EASY-Spray nanoelectrospray ionization source (Thermo Fisher Scientific, Bremen, Germany). Protein digests were resolved on a trap column (C18 2 cm × 100 μm, 120 Å, SunChrom) at a flow rate of 10 μl/min using 0.1% formic acid and separated on an analytical column (25 cm × 150 μm, C18 1.9 μm, 120 Å, SunChrom) with a flow rate of 600 nl/min over a linear gradient from 5 to 40% solvent B (100% ACN, 0.1% formic acid) for 40 min. All columns were equipped in a column oven at 60 °C. The source was operated in the positive ion mode at 2.3 kV with a capillary temperature of 350 °C. The Orbitrap Exploris 480 was operated in the data-dependent mode at target mass resolutions of 60 000 and 15 000 (at m/z 400). The scan range was m/z 350-400, an intensity above the 50000 threshold with a charge ≥2 was selected, and fragmentation was performed by high-energy collision-induced dissociation with a normalized collision energy of 28. The maximum injection times for the survey scan and the MS/MS scan were 22 ms and 22 ms, respectively. To minimize the redundant selection of precursor ions, the fragmented ions were dynamically excluded for 30 s. The mass spectrometry proteomics dataset was deposited into Proteome Discoverer 2.4. The raw data were processed using Proteome Discoverer (version 2.4.1.15, Thermo Scientific). The fragmentation spectra were searched with the SEQUEST engine against the UniProtKB/Swiss-Prot Rat database (UniProt_Rattus_CON_20210415.fasta, release 2021_0415) using the following parameters: trypsin, maximum 2 missed cleavages, precursor mass tolerance of 10 ppm, fragment mass tolerance of 0.02 Da, cysteine carbamidomethylation as a fixed modification, and oxidation of methionine and acetylation of the N-terminus as dynamic modifications. The peptide spectral matches were filtered at a 1% false discovery rate using Percolator. The peptide abundance was estimated by extracting the areas of the precursor ions with Proteome Discoverer software. The protein abundance was estimated by summing the abundances of all unique peptides (14).

### Liquid chromatography-electrospray ionization mass spectrometry (LC-ESI-MS) assay of MUPs

All analyses were performed with a Q-TOF mass spectrometer (Agilent 6530, USA), fitted with an AJS ESI ion source and connected to a liquid chromatograph (Agilent 1290, USA). Two urine samples from six males of each subspecies were mixed in equal amounts to obtain three test samples. The urine samples were desalted and concentrated on a 300SB-C8 column (2.1× 50 mm, C8 3.5 μm, 300 Å, ZORBAX), and the proteins were eluted at a flow rate of 200 μl/min using three repeated 0-100% (v/v) acetonitrile (ACN) gradients. Data were collected between 600 and 2400 T (m/z), processed and transformed to a neutral average mass using Qualitative Analysis B.06.00 (MassHunter Workstation Software, Agilent) (13). ESI-MS allowed semiquantitative assessment of the relative amount of each MUP. The complete quality of *Mups* matched the mature protein, which was removed from the predicted signal peptide and subtracted from 2 Da for formation of a single disulfide bond predicted by the genome or cDNA sequence. For a particular protein, the intensity was quantified according to the peak height, and the relative intensity was a percentage of the MUP13 peak height (5).

### RNA-seq analysis of the VNO

The reference genome (rn6) index was built using Bowtie version 2.2.3, and paired-end clean reads were aligned to the reference genome using TopHat version 2.0.12. HTSeq version 0.6.1 was used to count the reads. Differentially expressed genes (DEGs) were identified using the DESeq1 R package version 1.24.0. An adjusted *P* < 0.05 and a |log_2_ fold change| > 1 were used as the criteria for significance (48). VNO transcriptome data of 9 females of each subspecies were downloaded from the NCBI database (BioProject ID: PRJNA591253).

### Construction of an eukaryotic expression vector

Rat *Vom2r53, Gαo, Gαi2* and *Trpc2* cDNAs were amplified by PCR from female rat VNOs (PrimeSTAR^®^ GXL DNA Polymerase Takara, Japan). *Vom1r68* was synthesized by BGI (China). The oligonucleotide primer sequences are shown in Supplementary Table S1. pME18S-Rho-*Vom1r68*, pME18S-*Rho*-*Vom2r53*, pME18S-*Gαo*, pME18S-*Gαi2* and pME18S-*Trpc2* were constructed with the Trelief^™^ SoSoo Cloning Kit (version 2.3, Tsingke, China) through homologous recombination. The constructed plasmid was transformed into DH5α cells (CB101, Tiangen, China). The positive clones were selected and sequenced using universal primers (T7 promoter and SP6 terminator primer). The strains were expanded and cultured for endotoxin-free plasmid extraction with the EndoFree Maxi Plasmid Kit V2 (DP120, Tiangen, China). The constructed plasmids were stored at −20 °C for the transfection of HEK293-T cells (49).

### Transfection of HEK293-T cells

HEK293-T cells were grown in DMEM supplemented with 10% foetal bovine serum (TBD 31HB, TBD sciences, China). The cells were maintained at 37 °C in a humidified atmosphere containing 5% CO_2_. The cells were seeded onto 24×24-mm sterile slides coated with 100 μg/ml poly-D-lysine. One hour before transfection, the fresh complete culture medium was replaced, and culture was continued at 37 *°C* in a humidified atmosphere containing 5% CO_2_. A total of 40-60% confluent cells were transfected with 2.5 μg of pME18S-*Rho*-*Vom1r68* (pME18S-*Rho*-*Vom2r53*), 1.5 μg of pME18S-*Gαi2* (pME18S-*Gαo*) and 1.0 μg of pME18S-*Trpc2* using VigoFect (Vigorous Biotechnology, China) (2.5 μl/μg DNA). A signal peptide (RHO) was added to the N-terminus of the vomeronasal receptor to facilitate vomeronasal receptor expression on the membrane (50). After 24 h of incubation, the Fura-2-AM (Beyotime, S1052, China) fluorescent probe was incubated for 40 min at 37 °C (49).

### Immunofluorescence

HEK293-T cells were split and seeded on poly-D-lysine-coated glass coverslips. The cells were then transfected with 2.5 μg of pME18S-*Rho*-*Vom1r68* (pME18S-*Rho*-*Vom2r53*), 1.5 μg of pME18S-*Gαi2* (pME18S-*Gαo*), 1.0 μg of pME18S-*Trpc2* and 1.0 μg of mTmG, which could be expressed on the cell membrane using VigoFect (Vigorous Biotechnology, China) (2.5 μl/μg DNA) (51). After 48 h, the HEK293-T cells were washed three times with PBS, 4% paraformaldehyde in PBS pH 7.4 was added, and the cells were maintained for 10 min at room temperature. HEK293-T cells were washed three times with PBS and incubated with 1% BSA and 22.52 mg/mL glycine in PBST for 1 h at 37 °C. Primary incubation was conducted with mouse monoclonal anti-rhodopsin antibody 4D2 (ab98887 Abcam) (diluted 1:100 in 3% BSA) in a humidified chamber overnight at 4 °C. The solution was decanted, and the cells were subjected to three 5-min washes with PBS. The cells were incubated with an Alexa 488-conjugated donkey anti-mouse antibody (ab150105 Abcam) in 1% BSA for 1 h at room temperature in the dark. The secondary antibody solution was decanted and subjected to three 5-min washes with PBS in the dark. The cells were incubated with 1 μg/mL Hoechst stain for 1 min and rinsed with PBS. The coverslip was mounted with a drop of mounting medium and sealed with nail polish. Images were acquired using a confocal laser microscope (Zeiss LSM 710) (52, 53).

### Preparation of 2-heptanone and recombinant MUP13

2-Heptanone (purity > 98%) was purchased from ACROS Organics. Ringer’s solution (140 mM NaCl, 5 mM KCl, 1 mM CaCl_2_, 1 mM MgCl2, 1 mM sodium pyruvate, 10 mM glucose and 10 mM HEPES, pH 7.2) and High-K Ringer’s solution (40 mM NaCl, 100 mM KCl, 1 mM CaCl_2_, 1 mM MgCl_2_, 1 mM sodium pyruvate, 10 mM glucose and 10 mM HEPES, pH 7.2) were prepared.

The coding sequence of MUP13 (UniProt ID: P02761) was amplified by PCR using the primers F/ex/*Nde*I (5’-GGGTTTCATATGCATGCAGAAGAAGCTAGTTCCACAAGAG-3’) and R/*ex*/*XhoI* (5’-CCGCTCGAGTCCTCGGGCCTGGAGACAG-3’). The amplicon was double digested with *NdeI* and *Xho*I, cloned into the corresponding site of the pET28a expression vector (Novagen, Madison, WI, USA), transformed into *Escherichia coli* Trans1-T1 competent cells (TransGen, Beijing, China), and then cultured in LB medium containing kanamycin. The plasmids were extracted and sequenced, and the confirmed pET28a-MUP13 plasmid was transformed into *E. coli* Rosetta 2 competent cells (Novagen, San Diego, CA, USA) and cultured in the medium. The plasmids were extracted and sequenced, and the confirmed transformants were cultured in LB medium containing kanamycin. IPTG was added to induce protein expression. The cells were lysed, and the recombinant proteins were purified using HisPur Cobalt Superflow Agarose (Thermo Scientific, Waltham, MA, USA), dialyzed against 1× phosphate-buffered saline (PBS) and then stored at −80 °C until use. SDS–PAGE and nLC–MS/MS (Thermo Scientific, Waltham, MA, USA) were used to assess the purification of rMUP13 and verify the identity of the recombinant protein, respectively (14, 22).

### Calcium imaging following transfection and exposure to pheromone analogues

The samples were illuminated with an ion imaging system (Nikon Ti-E, Japan), and the changes in the calcium ratio were monitored. Compounds were delivered at a flow rate of 1 ml/min by a peristaltic pump. At the end of each imaging session, 100 mM KCl in Ringer’s solution was applied to HEK293-T cells expressing vomeronasal receptor genes to check the viability and responsiveness of HEK293-T cells. The duration of the stimulus delivery was 2 min. The interstimulus intervals were 4 min or longer to allow the recovery of HEK293-T cells. Solutions of 2-heptanone, 4-heptanone and squalene were prepared as stock solutions of 1 M by dissolution in dimethyl sulfoxide (DMSO) and diluted in Ringer’s solution to 10^-5^ M. The solution of 10^-7^ M rMUP13 was a mixture of 30 μl of purified rMUP13 protein and 40 ml of Ringer’s solution, and the solution of 10^-7^ M rOBP3 was a mixture of 30 μl of purified rOBP3 protein and 40 ml of Ringer’s (20).

### Data analysis

We used the Kolmogorov–Smirnov test to examine the distribution of raw data and used either nonparametric tests or parametric tests for the subsequent analyses. Independent t tests or Mann–Whitney U tests were used to compare the abundance of urinary volatiles between the RNH and RNC subspecies. Independent-sample t tests were used to assess the differences in the expression of hepatic mRNA and urinary MUP VNO mRNA levels between the two subspecies. Independent-sample t tests and one-way ANOVA were used to determine the differences in the hepatic mRNA expression levels between the two subspecies. All statistical analyses were conducted using SPSS (Version 18.0). Significance was set to *P* < 0.05.

## Results

### Urinary volatile pheromones in RNH and RNC

We characterized 19 compounds from voided urine, including eight ketones, one sulfone and four phenols (Fig. 1A, B). Higher levels of seven ketones (3-ethyl-2-pentanone, 4-heptanone, 2-heptanone, 6-methyl-5-hepten-2-one, 3-ethyl-2,4-heptanedione, 9-hydroxy-2-nonanone and two isomers of dialkyl tetrahydro-2H-pyran-2-one) and lower levels of 9,12-octadecadienoic acid were detected in RNH males compared with RNC males (Fig. 1A, B). Other detected urinary volatile compounds, such as dimethyl sulfone, phenol, 4-methyl-phenol, 4-ethyl-phenol and squalene, did not differ between these two subspecies. In particular, 2-heptanone was the most abundant of all the volatile pheromones in both RNH and RNC males.

**Figure 1.**
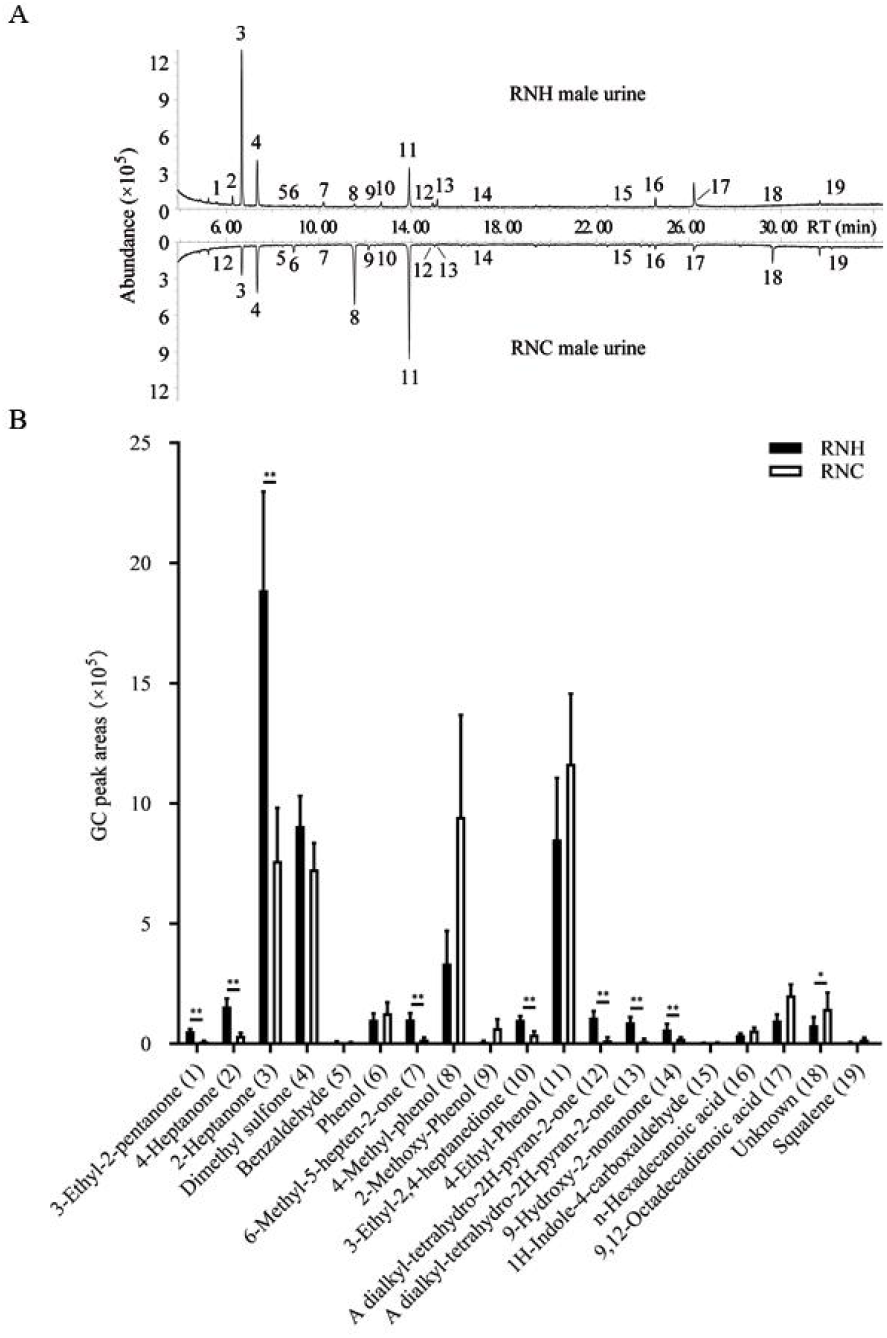
GC–MS assay of urinary volatile compounds in two rat subspecies. (A) Representative GC profile of dichloromethane extracts from the urine of male RNH and RNC rats. (B) Pairwise comparison of abundances quantified based on the GC peak areas of urinary volatiles between two subspecies (mean ± SE, n = 22 for RNH, n = 21 for RNC, \**P* < 0.05, \*\**P* < 0.01, independent-sample t test).

### Expression of hepatic *Mup* genes

The results from absolute quantitative PCR of the six known *Mup* genes detected in the liver showed that the expression of *Mup13* (t = 6.264, *P* = 0.002, n = 6), *Obp3* (t = 3.321, *P* = 0.020, n = 6) and alpha-2u globulin *Pgcl2* (t = 2.566, *P* = 0.028, n = 6) was significantly higher in RNH males than in RNC males. The expression of *Mup4* (t = 2.343, *P* = 0.060, n = 6, marginal significance) and *Mup5* (t = 2.040, *P* = 0.069, n = 6, marginal significance) tended to be higher in RNH males than in RNC males. No significant difference in the expression of *Pgcl3* (t = 0.533, *P* = 0.606, n = 6) was detected between RNH males and RNC males (Fig. 2A).

**Figure 2.**
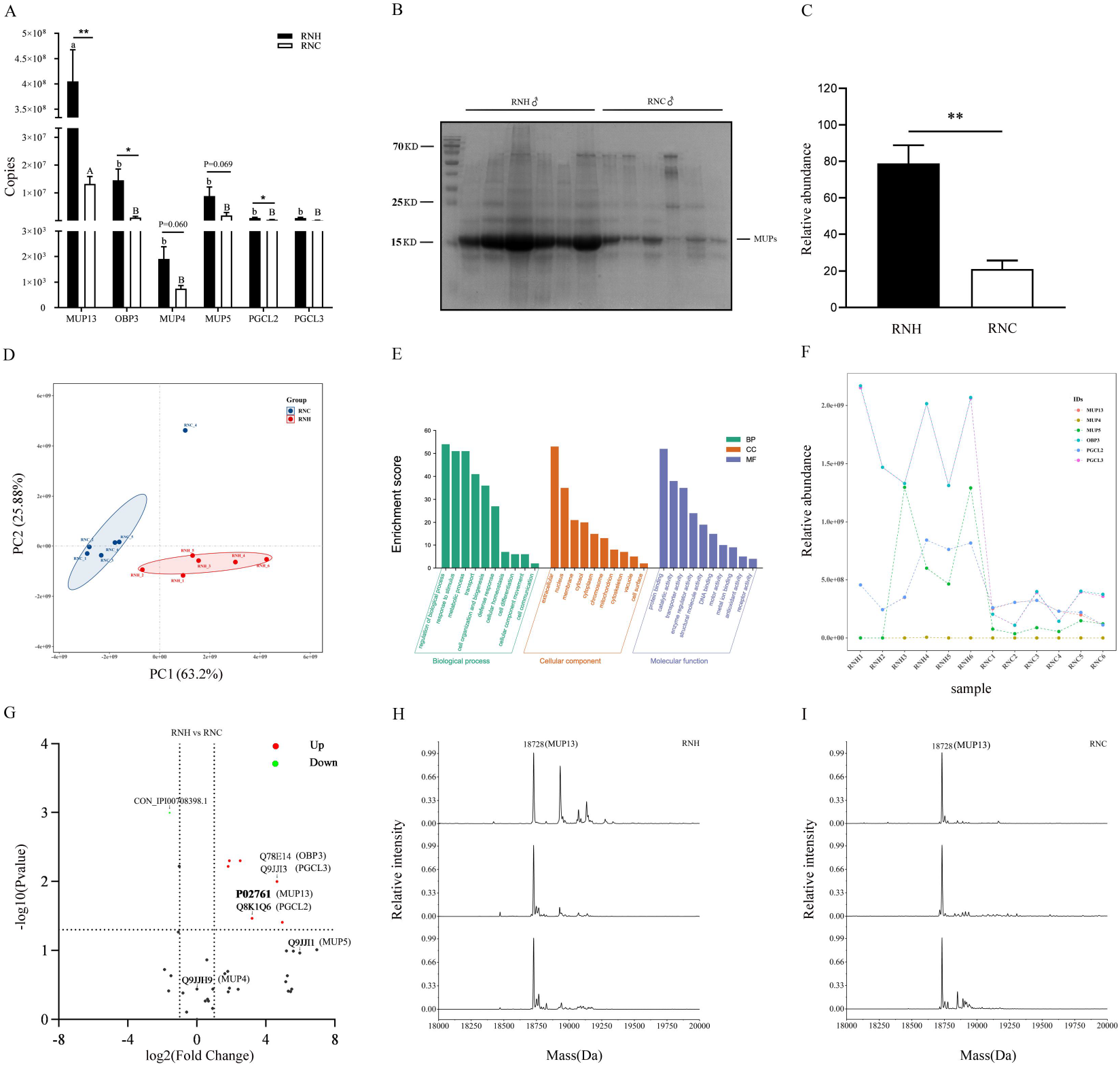
Hepatic *Mup* gene expression and MUP13 levels in urine in the two subspecies. A) Differences in the expression of hepatic *Mup* genes between RNH and RNC males [n = 6 for each subspecies, different lowercase letters indicate significant differences in expression between different hepatic Mup genes within the same subspecies (either RNH or RNC rats)]. * indicates a significant difference in the expression of the same hepatic gene between subspecies. (B) SDS–PAGE gel image of RNH and RNC male urine samples. (C) Relative abundance of MUPs in the urine of RNH and RNC males quantified by SDS–PAGE (n = 6 for each subspecies). (D) Plots of principal component 1 versus principal component 2 (PC1 vs. PC2) based on the expression of MUPs in RNH and RNC males. (E) Gene ontology enrichment analysis of genes in male urine showing expression differences between RNH and RNC subspecies. (F) Urinary relative abundance of MUPs in RNH and RNC males quantified by nLC–MS/MS (n = 6 for each subspecies). (G) Difference in male MUP13 expression between the two subspecies analysed by nLC–MS/MS. (H) The intact protein mass profiles of male urine of RNH and (I) RNC were analysed by LC-ESI-MS to obtain the profile of protein masses by focusing on the range of 18,000 Da to 20,000 Da (n = 3 for each subspecies). (mean ± SE, \**P* < 0.05, \*\**P* <0.01, independent-sample t test and one-way ANOVA).

The expression of the hepatic *Mup13* gene was highest among all identified *Mup* genes in both RNH and RNC males (Fig. 2A).

### Urine-borne MUP pheromones and their states

The RNH males had significantly higher MUP levels than the RNC males, as revealed by SDS–PAGE (t = 5.272, *P* = 0.000, n = 6) (Fig. 2B, C). Through in-gel digestion followed by nLC–MS/MS, we detected six MUPs [MUP13 (P02761), OBP3 (Q78E14), MUP4 (Q9JJH9), MUP5 (Q9JJI1), PGCL2 (Q8K1Q6) and PGCL3 (Q9JJI3)] in the urine of male RNH and RNC rats. A principal component analysis (PCA) of proteins in the urine of RNH and RNC males showed that they could be separated by PC1 (Fig. 2D). A Gene Ontology (GO) analysis of the male urine proteome data of the RNH and RNC subspecies showed that 41 transport proteins, including MUPs, were enriched in the biological process (BP) category (Fig. 2E). The whole protein sequence was used as an ExPASy entry, and the sequence coverage was 50%. MUP13 contained 181 amino acids, and the identified proteins shared similar molecular masses. The pI calculated by ExPASy-Compute pI/Mw was 6.21 for MUP13 (Table 1). The abundance of MUP13 (t = 3.074, *P* = 0.022, n = 6), OBP3 (t = 4.038, *P* = 0.010, n = 6), PGCL2 (t = 2.842, *P* = 0.034, n = 6) and PGCL3 (t = 4.047, *P* = 0.010, n = 6) in RNH males was significantly higher than that in RNC males. No significant difference in the abundance of MUP4 (t = 1.000, *P* = 0.363, n = 6) or MUP5 (t = 1.952, *P* = 0.108, n = 6) was detected (Fig. 2F, G). The ESI-MS analysis of the urine of RNH and RNC males revealed that MUPs dominated the deconvoluted mass spectra, and several proteins and multiple discrete masses in the 18-20-kDa mass range were evident. The ESI-MS results showed that the protein at 18728 (MUP13) presented the highest intensity in the urine of males of the two subspecies in all instances (Fig. 2H, I).

**Table 1.**
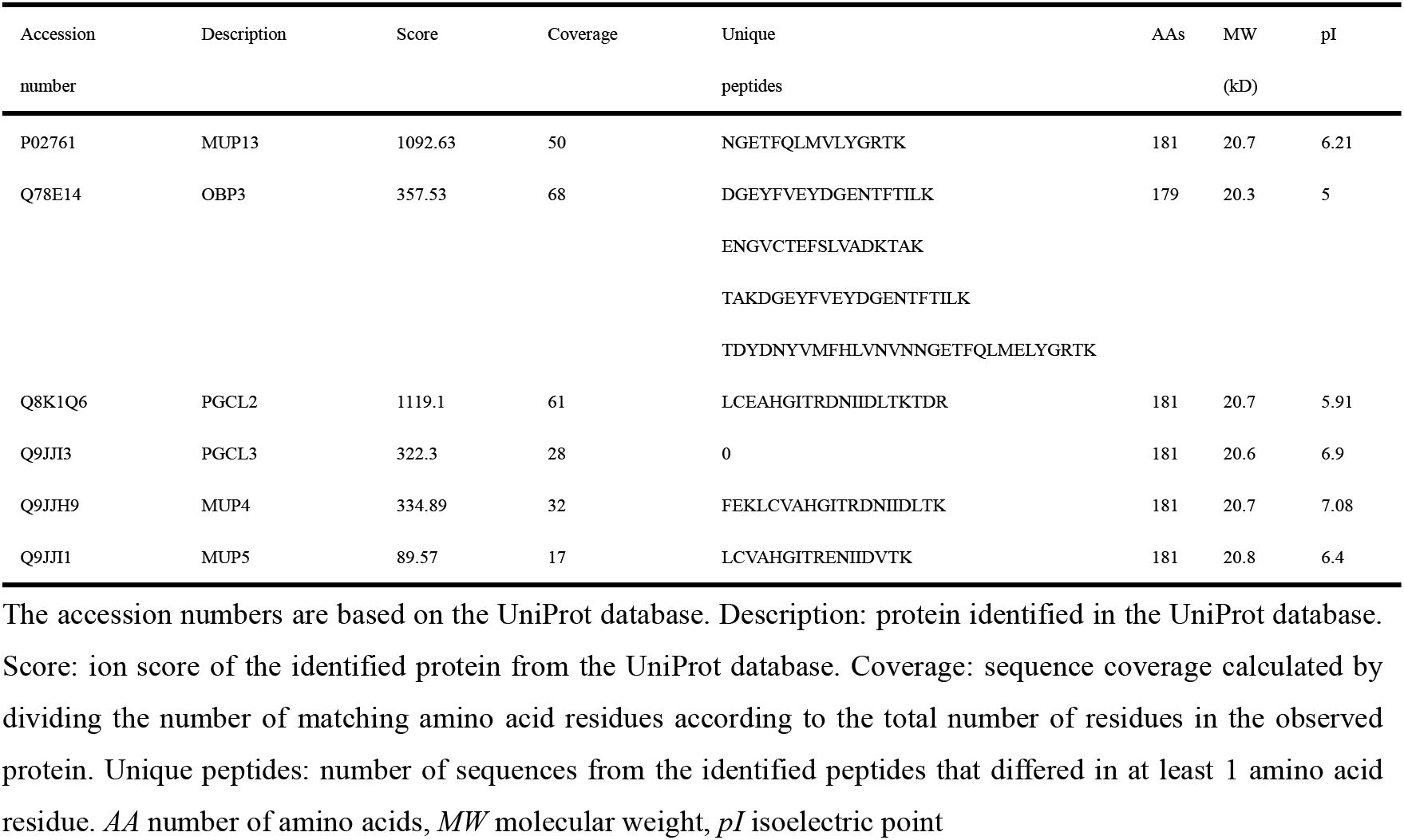
List of MUPs identified in male urine by nLC-MS/MS

### Transcriptome analysis of vomeronasal receptors

The MF-based Gene Ontology (GO) analysis of VNO transcriptome data from RNH and RNC females showed that 263 genes were enriched in the response to stimulus, including three *V1r* genes (*Vom1r60*, *Vom1r68* and *Vom1r81*) and three *V2r* genes (*Vom2r53*, *Vom2r-ps1* and *Vom2r43*). Gene Ontology (GO) enrichment analysis in terms of MF revealed 93 genes involved in the response to organic substances, including three V1Rs (*Vom1r60*, *Vom1r68* and *Vom1r81*) (Fig. 3A). In RNH and RNC, a total of 180 VRs were detected, including 93 V1Rs and 87 V2Rs. Three differentially expressed genes in the *V1r* family and three differentially expressed genes (including a pseudogene) in the *V2r* family were identified. In the *V1r* family, the expression level of *Vom1r68* was significantly higher in RNH females than in RNC females, and the expression levels of *Vom1r60* and *Vom1r81* were significantly lower in RNH females than in RNC females. In the *V2r* family, the expression levels of *Vom2r53* and *Vom2r pseudogene 1* were significantly higher in RNH females than in RNC females, and the expression level of *Vom2r43* was significantly lower in RNH females than in RNC females (Fig. 3B, C, E, F).

**Figure 3.**
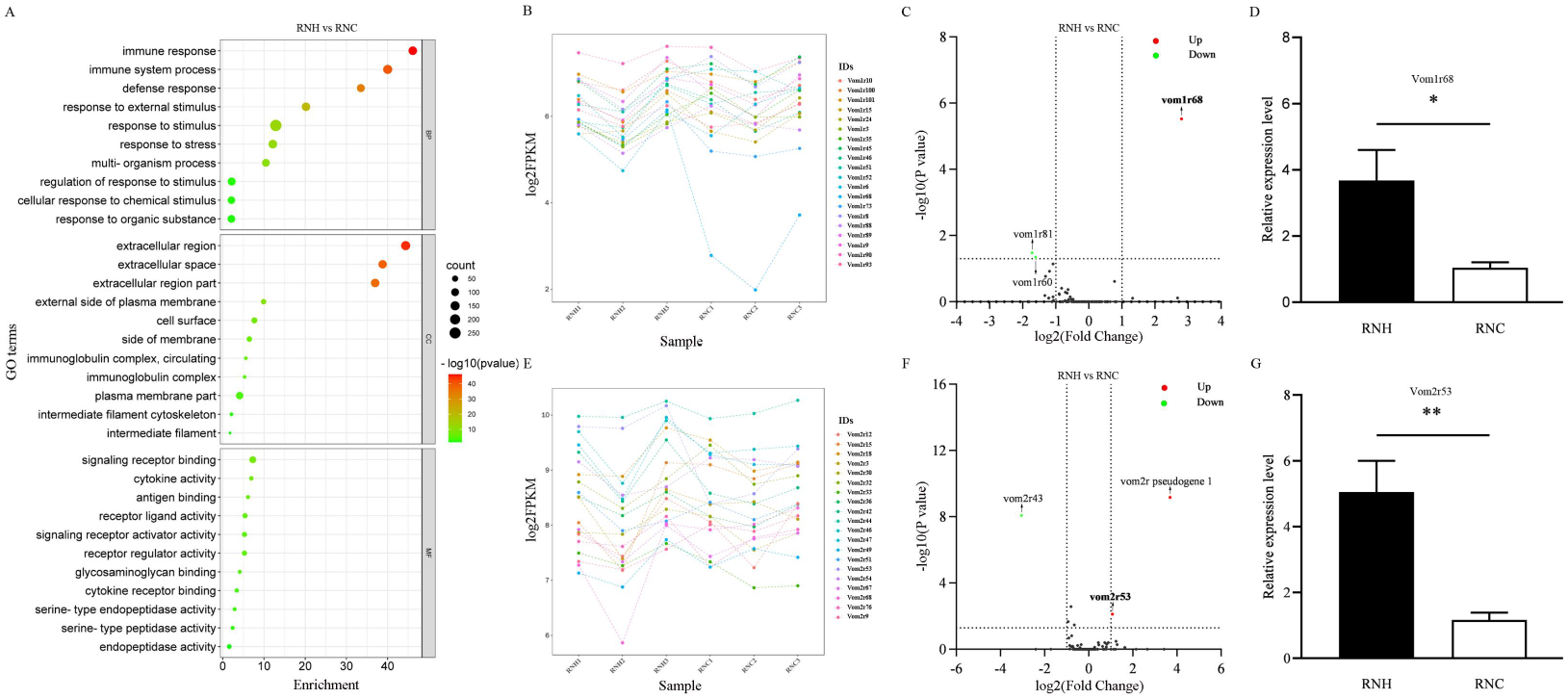
Candidate pheromone receptor genes. (A) Gene ontology enrichment analysis of differentially expressed vomeronasal receptor genes between females of the RNH and RNC subspecies. (B) Expression of the *V1r* family and (E) *V2r* family in females of the RNH and RNC subspecies revealed by transcriptome analysis. (C) Differences in the expression of the *V1r* family and (F) *V2r* family in females of the RNH and RNC subspecies revealed by transcriptome analysis. (D) Differences in the expression of *Vom1r68* (n = 4 for each subspecies) and (G) *Vom2r53* in females of each subspecies (n = 9 for each subspecies) revealed by qPCR-based comparison between the RNH and RNC subspecies (mean ± SE, \**P* < 0.05, \*\**P* <0.01, independent-sample t test).

### mRNA expression levels of *V1r* and *V2r* in RNH and RNC females

The qPCR results showed that the mRNA expression levels of *Vom1r68* (t = 2.811, *P* = 0.031, n = 4) and *Vom2r53* (t = 4.006, *P* = 0.003, n = 9) were significantly higher in RNH females than in RNC females (Fig. 3D, G).

### Immunofluorescence and calcium imaging

The immunofluorescence results showed that the Vom1r68 receptor colocalized with mTmG on the membrane of HEK293-T cells (Fig. 4A-C). A total of 802 HEK293-T cells transfected with *Vom1r68* were analysed, and 2.74% of the HEK293-T cells expressed the Vom1r68 receptor on the membrane. Using calcium ion imaging, we investigated three structurally diverse volatile pheromones in recipient female rats: 2-heptanone, 4-heptanone and squalene. A total of 1077 HEK293-T cells expressing *Vom1r68* were analysed, and 31 of these cells were activated by 10^-5^ M 2-heptanone, corresponding to an activation rate of 2.88%. However, the cells were not activated by 10^-5^ M 4-heptanone or 10^-5^ M squalene (Fig. 4D-H).

**Figure 4.**
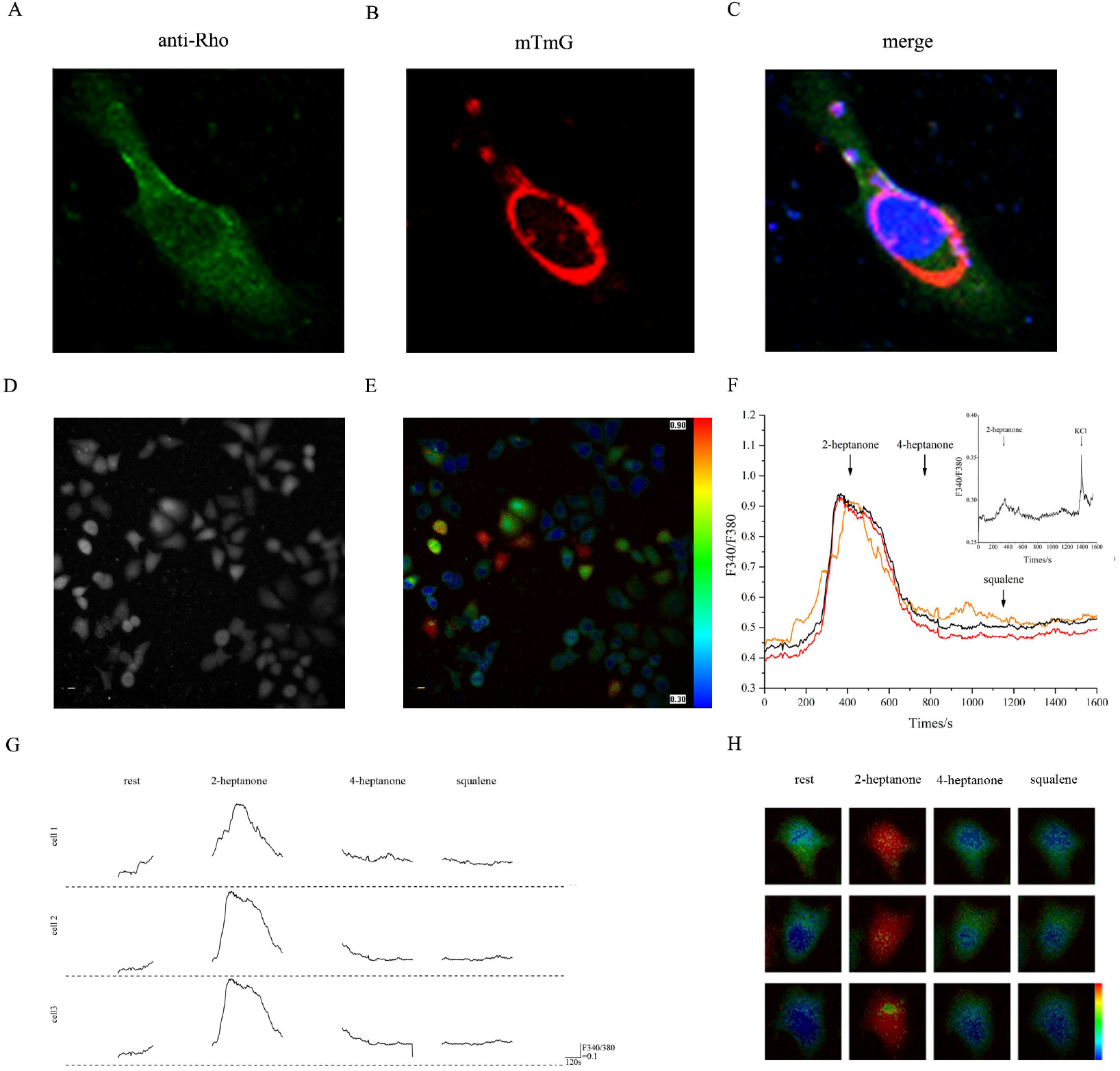
Pheromones that evoke intracellular Ca^2+^ elevation in single HEK293-T cells transfected with *Vom1r68*. (A-C) The Vom1r68 receptor colocalizes with mTmG on the membrane of HEK293-T cells. (D) Fluorescence images in grey and (E) pseudocolour scales. (F) Traces of three HEK293-T cells transfected with *Vom1r68* showing responsiveness to 10^-5^ M 2-heptanone and 100 mM KCl (positive control) but not to 10^-5^ M 4-heptanone and 10^-5^ M squalene. (G) Response profiles of three representative HEK293-T cells. (H) Corresponding Fura-2 ratio images of the three HEK293-T cells. Scale bar: 10 μm.

Immunofluorescence results showed that the Vom2r53 receptor colocalized with mTmG on the membrane of HEK293-T cells (Fig. 5A-C). A total of 908 HEK293-T cells transfected with *Vom2r53* were analysed, and 2.42% of the HEK293-T cells expressed the Vom2r53 receptor on the membrane. By calcium ion imaging, we investigated two MUP pheromones in recipient female rats: MUP13 and OBP3. A total of 1343 HEK293-T cells expressing *Vom2r53* were analysed, and 44 of these cells were activated by 10^-7^ M MUP13, corresponding to an activation rate of 3.28%. However, the cells were not activated by 10^-7^ M OBP3 (Fig. 5D-H).

**Figure 5.**
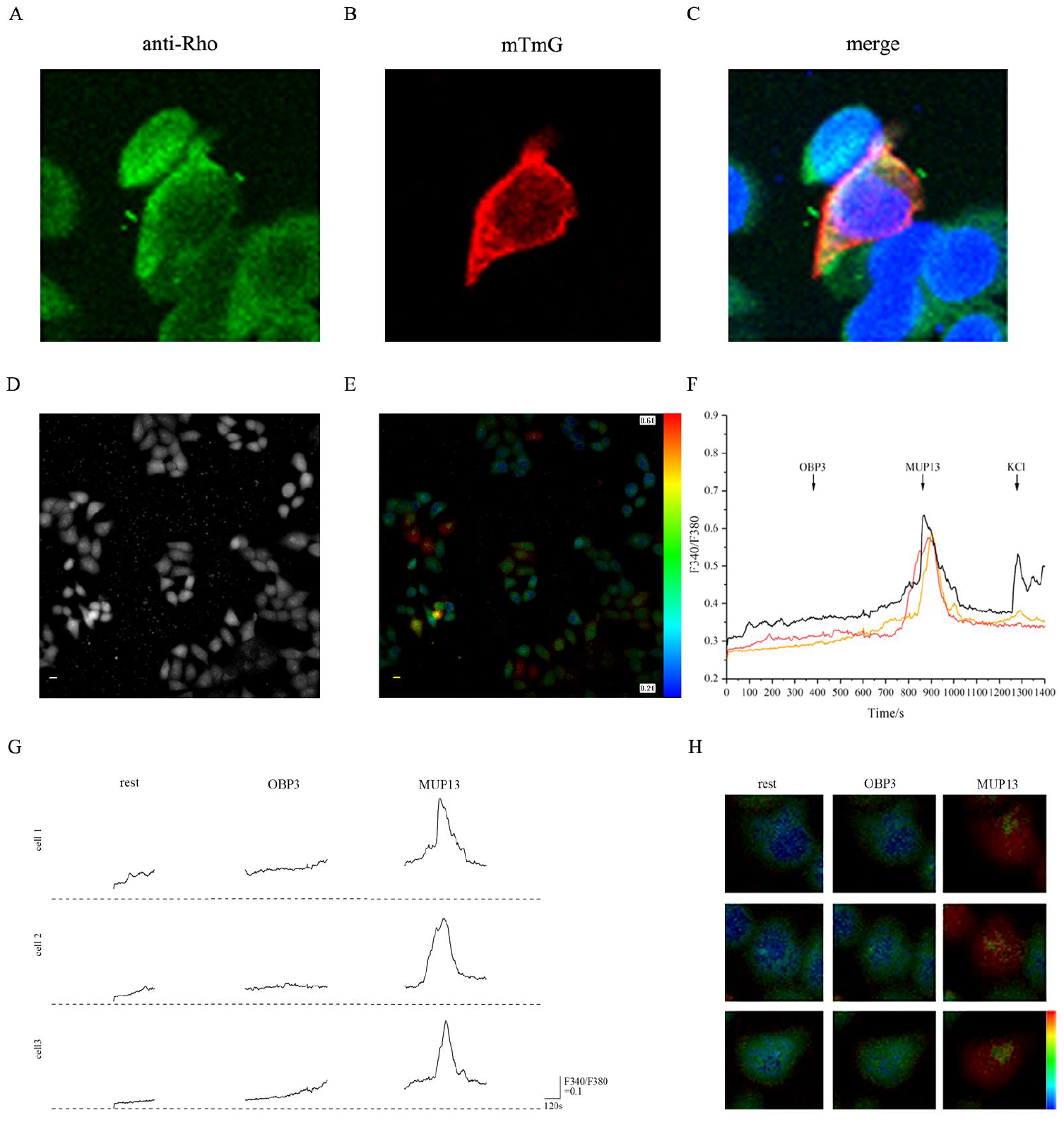
Pheromones that evoke intracellular Ca^2+^ elevation in single HEK293-T cells transfected with *Vom2r53*. (A) The Vom2r53 receptor colocalizes with mTmG on the membrane of HEK293-T cells. (D) Fluorescence images in grey and (E) pseudocolour scales. (F) Traces of three HEK293-T cells transfected with *Vom2r53* showing responsiveness to 10^-7^ M MUP13 and 100 mM KCl (positive control) but not to 10^-7^ M OBP3. (G) Response profiles of three representative HEK293-T cells. (H) Corresponding Fura-2 ratio images of the three HEK293-T-cell lines. Scale bar: 10 μm.

## Discussion

Our data showed that male RNH rats had higher levels of the male pheromones 2-heptanone and MUP13 than male RNC rats and accordingly showed higher expression of Vom1r68 and Vom2r53, which have been identified as the vomeronasal receptors of 2-heptanone and MUP13, respectively. We performed the first identification of two pheromone-specific receptors in rats and confirmed the coadaptation of pheromones and their receptors in mammals.

In the 30 years since mammalian olfactory receptors were first reported in rats, only a few mammalian pheromone receptors have been identified in mice but not rats (6, 7, 38). Because the first identification of sex pheromones occurred nearly 30 years later in rats than in mice, the study of pheromone receptors in rats was delayed (4, 10, 12, 14, 20, 21, 54, 55). In mice, well-known pheromones are usually the main components found in male urine and preputial glands (4, 11). As such, we confirmed that 2-heptanone and MUP13 were among the most abundant volatile compounds and MUP components in wild rats and found markedly higher concentrations in RNH rats than in RNC rats by integrated hepatic *Mup* gene expression analysis, urinary SDS–PAGE and gas/liquid chromatography–mass spectrometry approaches. In addition, we observed that the volatile pheromones 2-heptanone and MUPs showed sexual dimorphism and affected sex recognition in both RHN and RNC rats, which was representative of the characteristics of male pheromones in these two subspecies (Fig. S3, S4).

Pheromones and their receptors often show genetic coupling and coevolution (16, 18, 56). By recording the number of 2-heptanone- or MUP13-activated VSNs, we previously found that RNH females have more VSNs expressing 2-heptanone and MUP13 receptors than RNC females (20). In the current study, further transcriptome analysis confirmed that several vomeronasal receptor genes exhibited differential expression between RNH females and RNC females, and two of these genes (*Vom1r68* and *Vom2r53*) that were upregulated instead of downregulated in RNH females were speculated to be genetically associated with 2-heptanone and MUP13, whose levels were increased in RNH male rats. This speculation was further confirmed by heterologous expression of the receptor genes encoding Vom1r68, a 2-heptanone receptor, and Vom2r53, an MUP13 receptor, in HEK293-T cells.

The V1R family is specifically receptive to volatile pheromones (57). Here, we found that although 2-heptanone, a male pheromone, could activate HEK293-T cells expressing *Vom1r68*, 4-heptanone (an isomer of 2-heptanone and a minor male pheromone) failed to do so, indicating the specificity of *Vom1r68* as a 2-heptanone receptor. V1rb2 has been identified as the receptor corresponding to the volatile pheromone 2-heptanone in mice (6, 33, 58). In addition, an evolutionary analysis using the maximum likelihood method showed that the *Vom1r68* sequence of rats was highly similar to the *V1rb2* sequence of mice (Fig. S1, S2).

V2Rs have been shown to specifically receive nonvolatile pheromones such as MUPs, major histocompatibility complex (MHC) peptides, and ESP1 in mice (59). In rats, MUPs activate VSNs with V2Rs in the basal layer of VNOs (60). As the only predominant MUP component in rats, MUP13 might be the decisive MUP pheromone that activates VSNs (20). Here, we demonstrated that HEK293-T cells expressing *Vom2r53* could be activated by MUP13 instead of OBP3, a minor MUP pheromone, indicating that Vom2r53 is an MUP13-specific receptor.

Heterologous expression provides a convenient approach for conducting large-scale functional and deorphanization studies of G protein-coupled receptors (43). The ligands of pheromone receptors in insects have been successfully identified using heterologous expression systems, including *Xenopus* oocytes and the HEK293 cell line (61, 62). For example, olfactory pheromone receptors of moths and locusts that were identified by expression in *Xenopus* oocytes were confirmed through further genetic manipulations (45, 46). The identification of an olfactory receptor for a male pheromone has been realized using heterologous expression systems and gene manipulation (44). Additionally, a few known vomeronasal receptors for certain pheromones were initially identified using dissociated VSNs and calcium imaging in mutant mice rather than a heterologous system (6, 7). The results from the immunofluorescence analysis performed in the current study showed that these two VR receptors colocalized with mTmG on the membrane of HEK293-T cells, which suggested that VR receptors can be expressed on the membrane of HEK293-T cells despite the low expression efficiency. Therefore, the interaction between vomeronasal receptors and ligands can be verified using heterologous expression systems. The results of calcium ion imaging showed that 2.88-3.28% of the analysed HEK293-T cells expressing Vom1r68 or Vom2r53 were activated by either 2-heptanone or MUP13, consistent with previous results revealing low firing rates of neurons (63). The low proportion of HEK293-T cells expressing vomeronasal receptors that responded to pheromones may be related to the calreticulin protein blocking the expression of vomeronasal receptors on the membrane, as demonstrated in mice. Previous studies on the heterologous expression of vomeronasal receptors have mainly focused on mice rather than the rats used in the current study. Some differences in vomeronasal receptor genes, cofactors, and ion channels exist between these species (32). As a better model for studying human diseases, the rat is closer to humans in all aspects than the mouse and might be more suitable for the expression of vomeronasal receptors on the surface of HEK293-T cells (64). The current study constitutes the first attempt to study the interaction between pheromone receptors and ligands using HEK293-T cells in rats. Because 2-heptanone and MUP13 can activate the vomeronasal system of females and mediate female mate choice in rats, they should first activate Vom1r68 and Vom2r53, respectively, and the signals then project to the brain nucleus that regulates sexual behaviour (14, 20, 21, 54).

In conclusion, we first identified two vomeronasal receptors for two pheromones in rats and showed that the expression of these two receptors and their corresponding pheromone ligands was increased in RNH and decreased in RNC, indicating genetic coupling and coadaptation. This work might provide some results that differ from those found in mice and reveal two potential receptors that will serve as useful references for further work. The divergence of pheromones and their receptors between RNH and RNC rats is ultimately a coadaptation process, and these pheromones and receptors must remain coordinated in each subspecies for mating to occur (19). Signal reliability decreases with decreases in the intensity of the signal, and weak urine-borne pheromones might thus show dysfunctions in the modulation of mate choice in RNC rat populations (20, 65). Divergence in male pheromones and female receptors might contribute to evolving speciation; therefore, understanding the genetic architecture underlying pheromone-receptor coadaptation is essential for future research (19).

## Acknowledgements

This work was supported by grants from the Strategic Priority Research Program of the Chinese Academy of Sciences (Grant No. XDPB16 to JXZ) and the National Natural Science Foundation of China (Nos. 32070451 to YHZ and 31872227 to JXZ). pME18s and pME18s-RHO were kindly donated by Dr. Kazushige Touhara.

## Author Contributions

J.X.Z. and Y.H.Z. conceived the study. W.C.W., Y.Y.S., G.M.H. and Y.J.W. conducted heterologous expression; W.C.W. performed the MUP analysis and calcium imaging. Y.H.Z. and J.X.Z. analysed the volatile pheromones, performed the RNA-seq and screened out vomeronasal receptor candidates. W.C.W., Y.H.Z. and J.X.Z. wrote the manuscript. All the authors read and approved the final manuscript.

## Data availability

The RNA-seq datasets generated and/or analysed during the current study are available in the National Center for Biotechnology Information repository, [BioProject ID: PRJNA591253]. Other data generated or analysed during this study are included in this published article [and its supplementary information files].

## References

1. Brennan PA, Zufall F. Pheromonal communication in vertebrates. Nature. 2006;444(7117):308–15.

2. Wyatt TD. Chemical signals and signatures. Pheromones and Animal Behavior. New York: Cambridge University Press; 2014.

3. Brennan PA. Pheromones and mammalian behavior. In: Menini A, editor. The Neurobiology of Olfaction. Boca Raton, FL: CRC Press; 2010. p. 157–80.

4. Novotny MV. Pheromones, binding proteins and receptor responses in rodents. Biochemical Society Transactions. 2003;31:117–22.

5. Kimoto H, Haga S, Sato J, Touhara K. Sex-specific peptides from exocrine glands stimulate mouse vomeronasal sensory neurons. Nature. 2005;437(7060):898–901.

6. Boschat C, Pelofi C, Randin O, Roppolo D, Luscher C, Broillet MC, et al. Pheromone detection mediated by a *V1r* vomeronasal receptor. Nature Neuroscience. 2002;5(12):1261–2.

7. Haga S, Hattori T, Sato T, Sato K, Matsuda S, Kobayakawa R, et al. The male mouse pheromone ESP1 enhances female sexual receptive behaviour through a specific vomeronasal receptor. Nature. 2010;466(7302):118–22.

8. Chamero P, Marton TF, Logan DW, Flanagan K, Cruz JR, Saghatelian A, et al. Identification of protein pheromones that promote aggressive behaviour. Nature. 2007;450(7171):899–902.

9. Zhang JX, Liu YJ, Zhang JH, Sun LX. Dual role of preputial gland secretion and its major components in sex recognition of mice. Physiology & Behavior. 2008b;95(3):388–94.

10. Zhang JX, Sun LX, Zhang JH, Feng ZY. Sex- and gonad-affecting scent compounds and 3 male pheromones in the rat. Chemical Senses. 2008a;33(7):611–21.

11. Roberts SA, Simpson DM, Armstrong SD, Davidson AJ, Robertson DH, McLean L, et al. Darcin: a male pheromone that stimulates female memory and sexual attraction to an individual male’s odour. BMC Biology. 2010;8:75.

12. Zhang YH, Zhang JX. A male pheromone-mediated trade-off between female preferences for genetic compatibility and sexual attractiveness in rats. Frontiers in Zoology. 2014;11:1–11.

13. Gomez-Baena G, Armstrong S, Halstead J, Prescott M, Roberts S, McLean L, et al. Molecular complexity of the major urinary protein system of the Norway rat, *Rattus norvegicus*. Scientific Reports. 2019;9(1):10757.

14. Guo X, Guo H, Zhao L, Zhang YH, Zhang JX. Two predominant MUPs, OBP3 and MUP13, are male pheromones in rats. Frontiers in Zoology. 2019;15(1):6.

15. Boake CRB. Coevolution of senders and receivers of sexual signals: Genetic coupling and genetic correlations. Trends in Ecology & Evolution. 1991;6(7):225–7.

16. Andersson M, Simmons LW. Sexual selection and mate choice. Trends in Ecology & Evolution. 2006;21(6):296–302.

17. Fukamachi S, Kinoshita M, Aizawa K, Oda S, Meyer A, Mitani H. Dual control by a single gene of secondary sexual characters and mating preferences in medaka. BMC Biology. 2009;7:64.

18. Bousquet F, Nojima T, Houot B, Chauvel I, Chaudy S, Dupas S, et al. Expression of a desaturase gene, *desat1*, in neural and nonneural tissues separately affects perception and emission of sex pheromones in Drosophila. Proceedings of the National Academy of Sciences of the United States of America. 2012;109(1):249–54.

19. Xu M, Shaw KL. Genetic coupling of signal and preference facilitates sexual isolation during rapid speciation. Proceedings of the Royal Society B 2019;286(1913):20191607.

20. Zhang JX, Zhao L, Fu SH, Wang ZS, Zhang JX. Male pheromones and their reception by females are co-adapted to affect mating success in two subspecies of brown rats. Current Zoology. 2021;67(4):371–82.

21. Zhang YH, Tang MM, Guo X, Gao XR, Zhang JH, Zhang JX. Associative learning is necessary for airborne pheromones to activate sexual arousal-linked brain areas of female rats. Behavioral Ecology and Sociobiology. 2019;73(6):1–12.

22. Papes F, Logan DW, Stowers L. The vomeronasal organ mediates interspecies defensive behaviors through detection of protein pheromone homologs. Cell. 2010;141(4):692–703.

23. Beynon RJ, Armstrong SD, Claydon AJ, Davidson AJ, Eyers CE, Langridge JI, et al. Mass spectrometry for structural analysis and quantification of the Major Urinary Proteins of the house mouse. International Journal of Mass Spectrometry. 2015;391:146–56.

24. Mudge J, Armstrong S, McLaren K, Beynon R, Jl H, Nicholson C, et al. Dynamic instability of the major urinary protein gene family revealed by genomic and phenotypic comparisons between C57 and 129 strain mice. Genome Biology 2008;9(5):1–16.

25. Roberts SA, Prescott MC, Davidson AJ, McLean L, Beynon RJ, Hurst JL. Individual odour signatures that mice learn are shaped by involatile major urinary proteins (MUPs). BMC Biology. 2018;16(1):48.

26. Wu DL. Subspecies of the brown rat *Rattus norvegicus* Berkenhout in China. Acta Theriologica Sinica. 1982;2:107–12.

27. Musser GM, Carleton MD. Superfamily Muroidea. In: Wilson DE, Reeder DM, editors. Mammal species of the world: a taxonomic and geographic reference: Johns Hopkins University Press; 2005. p. 894–1531.

28. Teng HJ, Zhang YH, Shi CM, Mao FB, Cai WS, Lu L, et al. Population Genomics Reveals Speciation and Introgression between Brown Norway Rats and Their Sibling Species. Molecular Biology and Evolution. 2017;34(9):2214–28.

29. Chen Y, Zhao L, Teng H, Shi C, Liu Q, Zhang J, et al. Population genomics reveal rapid genetic differentiation in a recently invasive population of Rattus norvegicus. Frontiers Zoology. 2021;18(1):6.

30. Mucignat-Caretta C. The rodent accessory olfactory system. Journal of Comparative Physiology A-neuroethology Sensory Neural and Behavioral Physiology. 2010;196(10):767–77.

31. Isogai Y, Si S, Pont-Lezica L, Tan T, Kapoor V, Murthy VN, et al. Molecular organization of vomeronasal chemoreception. Nature. 2011;478(7368):241–5.

32. Tirindelli R. Coding of pheromones by vomeronasal receptors. Cell and Tissue Research 2021;383(1):367–86.

33. Leinders-Zufall T, Lane AP, Puche AC, Ma WD, Novotny MV, Shipley MT, et al. Ultrasensitive pheromone detection by mammalian vomeronasal neurons. Nature. 2000;405(6788):792–6.

34. Munger SD, Leinders-Zufall T, Zufall F. Subsystem organization of the mammalian sense of smell. Annual Review of Physiology. 2009;71:115–40.

35. Touhara K, Vosshall LB. Sensing odorants and pheromones with chemosensory receptors. Annual Review of Physiology. 2009;71:307–32.

36. Chamero P, Katsoulidou V, Hendrix P, Bufeaf B, Roberts R, Matsunamib H, et al. G protein G_ao_ is essential for vomeronasal function and aggressive behavior in mice. Proceedings of the National Academy of Sciences of the United States of America. 2011;108(31):12898–903.

37. Sam M, Vora S, Malnic B, Ma WD, Buck LB. Neuropharmacology. Odorants may arouse instinctive behaviours. Nature. 2001;412(6843):142.

38. Buck L, Axel R. A novel multigene family may encode odorant receptors: a molecular basis for odor recognition. Cell. 1991;65(1):175–87.

39. Dulac C, Axel R. A novel family of genes encoding putative pheromone receptors in mammals. Cell. 1995;83(2):195–206.

40. Herrada G, Dulac C. A novel family of putative pheromone receptors in mammals with a topographically organized and sexually dimorphic distribution. Cell. 1997;90(4):763–73.

41. Osakada T, Ishii KK, Mori H, Eguchi R, Ferrero DM, Yoshihara Y, et al. Sexual rejection via a vomeronasal receptor-triggered limbic circuit. Nature Communications. 2018;9(1):4463.

42. Von der Weid B, Rossier D, Lindup M, Tuberosa J, Widmer A, Col JD, et al. Large-scale transcriptional profiling of chemosensory neurons identifies receptor-ligand pairs in vivo. Nature Neuroscience. 2015;18(10):1455–63.

43. Dey S, Zhan S, Matsunami H. A protocol for heterologous expression and functional assay for mouse pheromone receptors. Methods in Molecular Biology 2013;1068:121–31.

44. Yoshikawa K, Nakagawa H, Mori N, Watanabe H, Touhara K. An unsaturated aliphatic alcohol as a natural ligand for a mouse odorant receptor. Nature Chemical Biology. 2013;9(3):160–2.

45. Yang K, Huang LQ, Ning C, Wang CZ. Two single-point mutations shift the ligand selectivity of a pheromone receptor between two closely related moth species. Elife. 2017;6.

46. Guo X, Yu Q, Chen D, Wei J, Yang P, Yu J, et al. 4-Vinylanisole is an aggregation pheromone in locusts. Nature. 2020;584(7822):584.

47. Bustin SA, Benes V, Garson JA, Hellemans J, Huggett J, Kubista M, et al. The MIQE guidelines: Minimum information for publication of quantitative real-time PCR experiments. Clinical Chemistry. 2009;55(4):611–22.

48. Ibarra-Soria X, Nakahara TS, Lilue J, Jiang Y, Logan DW. Variation in olfactory neuron repertoires is genetically controlled and environmentally modulated. Elife. 2017;6:e21476.

49. Yoshikawa K, Touhara K. Myr-Ric-8A enhances G(alpha15)-mediated Ca^2+^ response of vertebrate olfactory receptors. Chemical Senses. 2009;34(1):15–23.

50. Krautwurst D, Yau KW, Reed RR. Identification of ligands for olfactory receptors by functional expression of a receptor library. Cell. 1998;95(7):917–26.

51. Luo L, Wang Y, Li B, Xu L, Kamau PM, Zheng J, et al. Molecular basis for heat desensitization of TRPV1 ion channels. Nature Communications. 2019;10(1).

52. Dey S, Matsunami H. Calreticulin chaperones regulate functional expression of vomeronasal type 2 pheromone receptors. Proceedings of the National Academy of Sciences of the United States of America. 2011;108(40):16651–6.

53. Aizman O, Brismar H, Uhlén P, Zettergren E, Levey AI, Forssberg H, et al. Anatomical and physiological evidence for D1 and D2 dopamine receptor colocalization in neostriatal neurons. Nature Neuroscience. 2000;3(3):226–30.

54. Zhang YH, Zhao L, Guo X, Zhang JH, Zhang JX. Sex pheromone levels are associated with paternity rate in brown rats. Behavioral Ecology and Sociobiology. 2019;73(2):15.

55. Zhang YH, Zhang JX. Urine-derived key volatiles may signal genetic relatedness in male rats. Chemical Senses. 2011;36(2):125.

56. Charlton BD, Owen MA, Swaisgood RR. Coevolution of vocal signal characteristics and hearing sensitivity in forest mammals. Nature Communications. 2019;10(1):1–7.

57. Mombaerts P. Genes and ligands for odorant, vomeronasal and taste receptors. Nature Reviews Neuroscience. 2004;5(4):263–78.

58. Del Punta K, Leinders-Zufall T, Rodriguez I, Jukam D, Wysocki CJ, Ogawa S, et al. Deficient pheromone responses in mice lacking a cluster of vomeronasal receptor genes. Nature. 2002;419(6902):70–4.

59. Kazushige T. Molecular biology of peptide pheromone production and reception in mice. Advances in Genetics. 2007;59:147–71.

60. Krieger Jr, Schmitt A, Lo”bel D, Gudermann T, Schultz Gn, Breer H, et al. Selective activation of G protein subtypes in the vomeronasal organ upon stimulation with urine-derived compounds. Journal of Biological Chemistry. 1999;274(8):4655–62.

61. Große-Wilde E, Svatoš A, Krieger J. A pheromone-binding protein mediates the bombykol-induced activation of a pheromone receptor in vitro. Chemical senses. 2006;31(6):547–55.

62. Wetzel, Christian H, Behrendt, Hans-Jorg, Gisselmann, Gunter, et al. Functional expression and characterization of a Drosophila odorant receptor in a heterologous system. Proceedings of the National Academy of Sciences of the United States of America. 2001.

63. Kajiya K, Inaki K, Tanaka M, Haga T, Kataoka H, Touhara K. Molecular bases of odor discrimination: reconstitution of olfactory receptors that recognize overlapping sets of odorants. Journal of Neuroscience. 2001;21(16):6018–25.

64. Do Carmo S, Cuello AC. Modeling Alzheimer’s disease in transgenic rats. Molecular neurodegeneration. 2013;8(1):1–11.

65. Searcy WA, Nowicki S. Reliability and Deception in Signaling Systems. The Evolution of Animal Communication. Princeton: Princeton University Press; 2006.

