## Supplementary material for "Screening and identification of two pheromone receptors based on the coadaptation of pheromones and their receptors in rats": cnorm.ini

```
[targets]
base=69
masses=5
mass1=50
mass2=131
mass3=219
mass4=414
mass5=502
target1=1
target2=45
target3=55
target4=2.5
target5=2
[ATUNE.cnorm]
masses=5
69=513792
50=7505
131=282304
219=205376
414=63032
502=71864
[STUNE.cnorm]
masses=5
69=457984
50=4749
131=267008
219=213248
414=16832
502=13128
```
