## Supplementary material for "Screening and identification of two pheromone receptors based on the coadaptation of pheromones and their receptors in rats": PRE_POST.INI

```
[POSTRUN]
SOURCE=MS
SampleAmount=0
Multiplier=1
Application=G1701DA Version D.00.00.38  19-Nov-2001 Copyright � Agilent Technologies 1989-2001
Date=Thu Jul 16 14:18:29 2015
SmartCard=AGILENT TECHNOLOGIES,5973N,0,3.01.58
acqmode=0
TuneName=atune.u
TunePath=C:\MSDchem\1\5973N\
TuneDate= 3 Jul 2015   7:23 am
CImode=0
CIpolarity=0
SourceTemp=243
QuadTemp=150
Matrix=1
InitialAmt=1
FinalAmt=1
PrimaryFile=\data.ms
PrimaryType=hgfile
timestamp=MS ChemStation
[Scan Parameters]
scanstart1=0.00
lowmass1=30.00
highmass1=450.00
scanstart2=-1.00
lowmass2=50.00
highmass2=550.00
scanstart3=-1.00
lowmass3=50.00
highmass3=550.00
[Tune Values]
massgain=387
massoffset=2
amugain=2173
amuoffset=134
dcpolarity=0
emission=34.6102
electron=69.9216
emvolts=1435.29
entlensoffset=21.0824
entrance_lens=16
filament=1
ionfocus=90.1569
repeller=19.9035
width219=-0.0511765
[MS_Faults]
NUM_MS_Faults=0
[firmware]
MSSC=3.01.58
BSP=1.2/22
HTTP=0.1
LUI=01.158.20010814
[Sequence]
RunClass=1
Update_QI=0
Update_RF=0
Update_RT=0
RunType=Sample
Upd_RF=No Update
Upd_RT=No Update
Upd_QI=No Update
ALS_Bottle=38
Area% Report=Default
Lib. Search Rep.=Default
Quant Report=Default
Post-Quant Macro=Default
CR Database=Default
CR Spreadsheet=Default
_methpath$=C:\MSDCHEM\1\METHODS\
_methfile$=RAT2014.M
[SeqArgs]
SEQARGS_1=0
SEQARGS_2=0
SEQARGS_3=0
SEQARGS_4=0
SEQARGS_5=0
SEQARGS_6=0
SEQARGS_7=65535
SEQARGS_8=65535
[SeqArgs2]
SEQARGS_1=0
SEQARGS_2=0
SEQARGS_3=0
SEQARGS_4=0
SEQARGS_5=0
SEQARGS_6=0
SEQARGS_7=0
SEQARGS_8=0
[Type]
Type=1
```
