## Supplementary material for "Screening and identification of two pheromone receptors based on the coadaptation of pheromones and their receptors in rats": PRE_POST.INI

```
[POSTRUN]
SOURCE=MS
SampleAmount=0
Multiplier=1
Application=G1701DA Version D.00.00.38  19-Nov-2001 Copyright � Agilent Technologies 1989-2001
Date=Tue Jul 14 00:45:55 2015
SmartCard=AGILENT TECHNOLOGIES,5973N,0,3.01.58
acqmode=0
TuneName=atune.u
TunePath=C:\MSDchem\1\5973N\
TuneDate= 3 Jul 2015   7:23 am
PrimaryFile=\data.ms
PrimaryType=hgfile
timestamp=MS ChemStation
Matrix=1
InitialAmt=1
FinalAmt=1
[Scan Parameters]
scanstart1=0.00
lowmass1=30.00
highmass1=450.00
scanstart2=-1.00
lowmass2=50.00
highmass2=550.00
scanstart3=-1.00
lowmass3=50.00
highmass3=550.00
```
