## Supplementary material for "Screening and identification of two pheromone receptors based on the coadaptation of pheromones and their receptors in rats": Supplementary materials.pdf

**Figures and Tables**

**Table S1 List of primers**

| Primer | list |
|---|---|
| Vom2r53-F | ATGTTAGAATTGGCCCATGGCAC |
| Vom2r53-R | CTATGTTTCAGAATTTTTACTCC |
| Gao-F | ATGGGATGTACTCTGAGCGCAGAGG |
| Gao-R | TCAGTACAAGCCACAGCCCCGGAGAT |
| Gai2-F | ATGGGCTGCACCGTGA |
| Gai2-R | TCAGAAGAGGCCACAGT |
| Trpc2-F | ATGGATCCCCTTTCGCCCAACTGGACTG |
| Trpc2-R | TTAGGACTCGCCCTTGGTCTCCAGATCTTC |
| Mup13-F | GGGAACCTCGATGTGGCTAA |
| Mup13-R | GCACTCTCCATTTTCCTTAATACG |
| Obp3-F | GTTGCCGACAAAACAGCAAAG |
| Obp3-R | AAGGATGAAGAAATAGATCTTGCCC |
| Mup4-F | GACAGATATGTCATGATTCACCTTG |
| Mup4-R | GCCTTGAGACAGCGATCAGT |
| Mup5-F | CCAGGACTCCAGCATCAAC |
| Mup5-R | AGGGAAGACGCTGGAGAGAC |
| Pgcl2-F | ACGGCAGAACAAAGGATCTGA |
| Pgcl2-R | TCAGGTACCAGTTCCACGTTA |
| Pgcl3-F | CCACAACAGGGAACCGTGA |
| Pgcl3-R | GCTGCACAAAAACTCTCATGC |
| T7 | TAATACGACTCACTATAGGG |
| SP6 | ATTTAGGTGACACTATAG |
| Gapdh-F | GACAATGAATATGGCTACAGCAAC |
| Gapdh-R | TTTATTGATGGTATTCGAGAGAAGG |

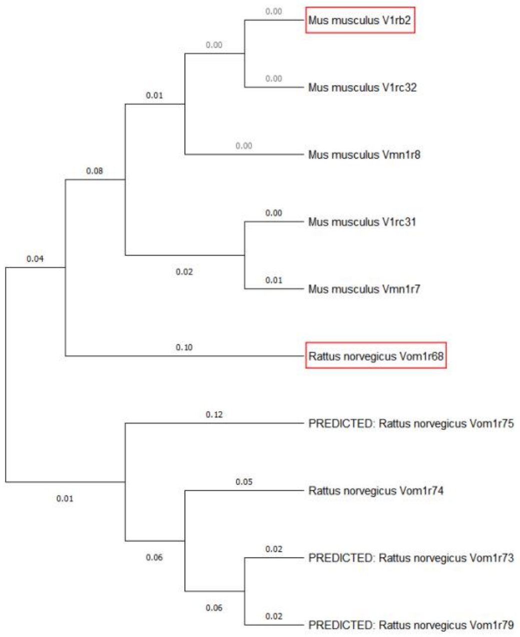


**Figure S1 Molecular evolutionary tree of rat and mouse *V1r* family**

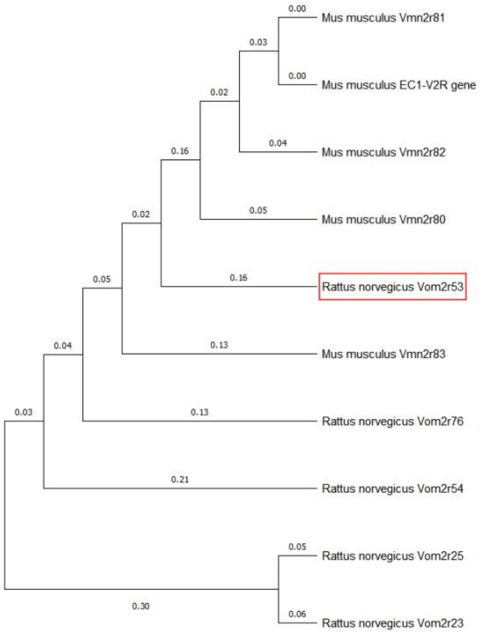


**Figure S2 Molecular evolutionary tree of rat and mouse *V2r* family**

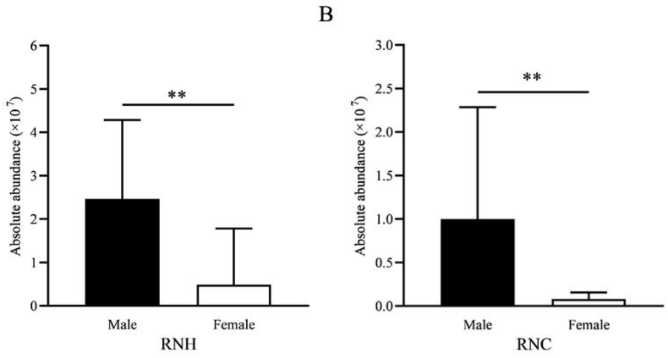


**Figure S3 The comparison of the abundance of 2-heptanone in male and female urine of RNH (n = 10) and RNC ( n = 9 for male, n = 10 for female ) (mean ± SE, ***$P$ < 0.01, independent-sample t-test).**

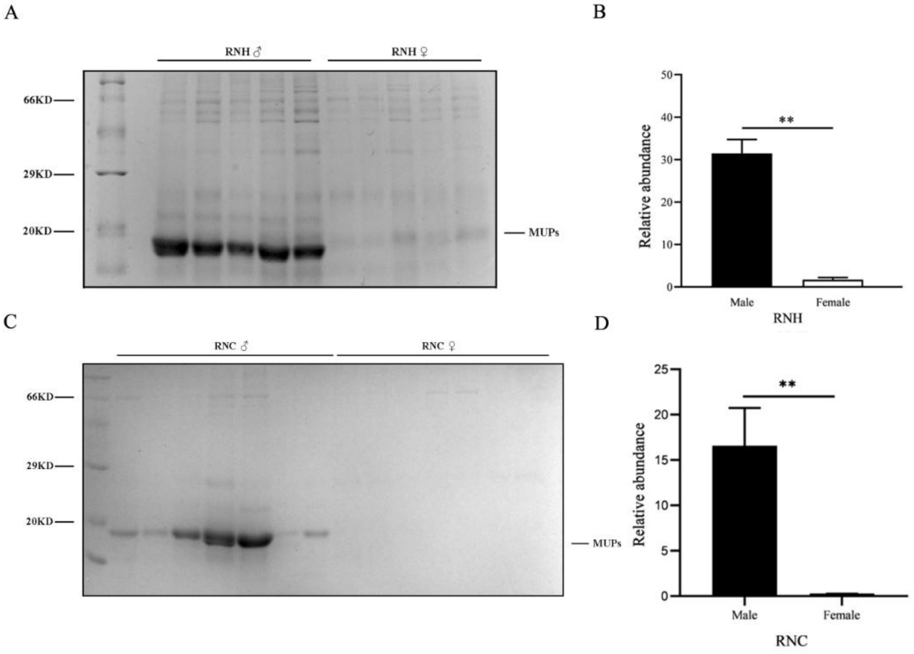


**Figure S4 SDS-PAGE gel image in male and female urine samples of RNH and RNC (n = 5, mean ± SE, ***$P$ < 0.01, independent-sample t-test).**
